# Absolute hand determination of glycofibrils from natural sources in cryo-EM

**DOI:** 10.1101/2025.09.30.679555

**Authors:** Qi Zhang, Lanju Qin, Tongtong Wang, Zhangqiang Li, Yilin Zhang, Sheng Chen, Nieng Yan, Jiawei Wang, Mingxu Hu

## Abstract

Glycans are one of the four fundamental macromolecules that constitute life. However, compared with proteins and nucleic acids, our understanding of the structures and functional mechanisms of glycans remains limited. Moreover, it is generally challenging to obtain high-resolution three-dimensional structures of glycans. Recent studies have demonstrated the potential of the CryoSeek strategy in enabling high-throughput structural determination of glycans, thereby presenting novel avenues for their structural investigation. Meanwhhile, unlike proteins, near atomic resolution density maps of glycofibrils do not inherently facilitate the determination of absolute hand, which is a prerequisite for building atomic models of glycofibrils. Existing absolute hand determination methods have severe limitations in the cases of glycofibrils from natural sources. In this study, we introduce Ahaha, a straightforward and efficient method for determining such absolute hand in cryo-EM. With their absolute hand measured by Ahaha, we built atomic models of four glycofibrils derived from a natural water sample, facilitating the study of glycans. The online service of Ahaha is available at https://cryoseek.org/ahaha.

## 1 Introduction

Glycans, alongside nucleic acids, proteins, and lipids, constitute the four fundamental classes of macromolecules that underpin the structural and functional essence of life. Beyond their well-established roles as energy sources and carbon precursors for biosynthesis, glycans mediate a diverse spectrum of biological functions, including acting as dynamic regulators of metabolism, cellular signaling, and intercellular communication [1]. These biological functions derive fundamentally from their unique structural properties [2–5]. The features of sugar molecules, including their high diversity, the presence of multiple chiral centers, and the possession of multiple sites capable of forming glycosidic linkages, result in the formation of highly complex glycan structures with diverse biological activities.

Due to their complicated stereochemistry and regiochemistry, the structural determination of glycans is formidable, thereby impeding our comprehensive understanding of the functions and mechanisms of glycans. A recently proposed research strategy, CryoSeek [6–8], which employs cryogenic electron microscopy (cryo-EM) for bioentity discovery, has revealed the widespread presence of glycoproteins and other glycoconjugates in the form of helical assemblies within natural environments [6–11]. We refer to these structures as “glycofibrils from natural sources”; for brevity, “glycofibrils” will be used hereinafter unless the “from natural sources” attribute needs specific emphasis. Notably, their inherent structural stability and helical assembly render them particularly amenable to high-resolution structure determination via cryo-EM. Therefore, the CryoSeek strategy paves the path for high-throughput structural determination of glycans.

However, cryo-EM transmission imaging intrinsically loses absolute hand information [12, 13]. Consequently, 3D reconstruction yields a density map that is randomly assigned to one of the two possible mirroring configurations (later referred as the enantiomeric pair). In the absence of folded protein sequences, near atomic resolution density maps of glycofibrils do not inherently enable the determination of absolute hand. For proteins, the prior knowledge that *α*-helices in proteins are exclusively right-handed can be leveraged to determine the absolute hand, specifically to decide whether the maps should be flipped. However, as demonstrated in our previous study of TLP-4 [7], many types of glycofibrils lack *α*-helices, as they are predominantly or purely composed of glycans, rendering such approaches inapplicable. Moreover, given that both D-sugars and L-sugars coexist in biological systems, even with ultra-high resolution (enabling the observation of individual carbon and oxygen atoms), it remains impossible to infer absolute hand based on the configurations of sugars. The ambiguity regarding the absolute hand of glycofibril density maps [7] impedes building atomic models from such maps and thereby obstructs structural investigation of glycans. Therefore, the absolute hand determination, or say, identifying which enantiomer in the enantiomeric pair is correct, requires addressing. We would like to emphasize that we deliberately choose the term “absolute hand” rather than “handedness” to avoid conceptual ambiguity regarding left-handedness or right-handedness of helices (see Supplementary Note I for further discussion).

Existing experimental methods for determining absolute hand are plagued by significant limitations when applied to glycofibrils. Tilt-pair imaging constitutes a classic approach for absolute hand determination [12–14], which images the same sample twice: first at a zero tilt angle and subsequently at a non-zero tilt angle (e.g. 20° [13, 14]). This method requires a pre-existing reference map to determine particle poses and thereby assign the absolute hand of the reconstruction. Meanwhile, the nature-sourced sample, taking the natural water as an example, comprises dozens, if not hundreds, of fibril types. Consequently, the requirement for tilt-pair imaging presents a fundamental barrier to the investigation of glycofibrils. Glycofibrils are characterized by high heterogeneity, and the identity of individual particles within this population remains undetermined prior to the attainment of high-resolution reconstruction. Given the limitations of the tilt-pair method under these conditions, its application to nature-sourced samples becomes prohibitively costly and labor-intensive (see Supplementary Note II for further discussion). Other methods include rotary shadowing [15], unidirectional shadowing [16], scanning electron microscopy, and atomic force microscopy [17]; these approaches require additional experimental procedures. Furthermore, their success is case-specific and highly unlikely to be applicable to thin glycofibrils, which typically possess a diameter in the range of 3 to 6 nanometers. Additionally, the instrumentation required for their implementation is not readily accessible in the majority of cryo-EM facilities. Finally, cryogenic electron tomography (cryo-ET) [18, 19], is also unlikely to be sufficient for determining the absolute hand of glycofibrils, despite the fact that the data it acquires contains absolute hand information, owing to its limited resolution and throughput. Given the absence of existing methods for determining the absolute hand of glycofibrils, a novel approach is thus required.

In this study, we propose Ahaha, shorthand for **A**bsolute **ha**nd determination of **h**elical **a**ssembly, a method for determining the absolute hand of glycofibrils from natural sources. In contrast to the conventional tilt-pair approach, Ahaha requires only mono-tilt micrographs, which can be readily acquired in a high-throughput manner in cryo-EM facilities. This manuscript is structured as follows. In Section 2.1, we outline the underlying principle of Ahaha. In Section 2.2, the correctness of absolute hand determination using Ahaha is validated with bacterial pili, which are also present in the dataset derived from a cave freshwater sample. In Section 2.3, we report the absolute hand measurements of four glycofibrils within the same dataset. Subsequently, in Section 2.4, atomic models of these four glycofibrils are built, enabling the structural investigation of glycans.

## 2 Results

### 2.1 Principle of Ahaha

In the research on glycofibrils, the timing of Ahaha application is as follows: in preceding experiments, viable density maps, albeit enantiomeric, are obtained, with helical parameters including rise (*r*) and twist (*θ* or 360° − *θ*) having been determined via helical indexing or other methods (Table 1).

**Table 1.** Bacterial pili and glycofibrils with their helical parameters in the studied water sample.

| fibril | type | rise (Å) | twist | twist (mirrored) |
| --- | --- | --- | --- | --- |
| pilus-like-α | bacterial pilus | 11.70 | 283.86° | 76.14° |
| pilus-like-β | bacterial pilus | 10.08 | 100.90° | 259.10° |
| pilus-like-γ | bacterial pilus | 10.40 | 112.27° | 247.73° |
| TLP-2a | glycofibril | 6.13 | 108.79° | 251.21° |
| TLP-2f | glycofibril | 5.54 | 132.05° | 227.95° |
| TLP-2g | glycofibril | 5.79 | 136.31° | 223.69° |
| TLP-2h | glycofibril | 6.14 | 93.78° | 266.22° |

The first step in the Ahaha workflow for absolute hand determination involves collecting a dataset consisting of mono-tilt micrographs. Empirically, the quantity of such micrographs is of the same order of magnitude as that required for CryoSeek (typically tens of thousands) when analyzing nature-sourced samples. The absolute hand measurement of Ahaha relies exclusively on mono-tilt micrographs, as opposed to the tilt-pair micrographs employed in the classical method. The theoretical basis of Ahaha is that helical assemblies, also referred to as fibrils (including glycofibrils), predominantly lie flat and parallel to the thin vitreous ice within the cryo-EM grid. This forms the prior knowledge concerning the pose distribution of particles extracted along the fibrils. In other words, these particles exhibit a pre-known pose distribution, where the tilt component of their poses should be congruent with the tilt of the vitrified ice (i.e., the tilt of the stage). Therefore, the axiom of Ahaha is to examine such congruence. As will be elaborated below, such tilt congruence only needs to hold at the level of sign, thereby rendering Ahaha robust.

The principle of Ahaha is illustrated as follows. The coordinate system conventions adopted in this study is depicted in Figure 1a, along with the diagram illustrates the optical path of the cryo-EM system with a lens configuration set to the underfocused condition. Particles observed in cryo-EM are Z-axis projections of the helices. As the stage tilts (for brevity, we adopt tilts of 20° or −20°, though any other angle is also applicable), the Z-axis projections of the two helical maps in the enantiomeric pair follow two rules. The first rule is that, for a helical map, tilting at 20° and −20° produces distinct Z-axis projections. This is because helical maps are assembled from asymmetric units. The Z-axis projections of an asymmetric unit differ when subjected to tilts of 20° and −20°. The second rule is that the Z-axis projection of one helical map when tilted at 20° is identical to the Z-axis projection of its mirroring counterpart in the enantiomeric pair when tilted at −20°.

**Fig. 1.**
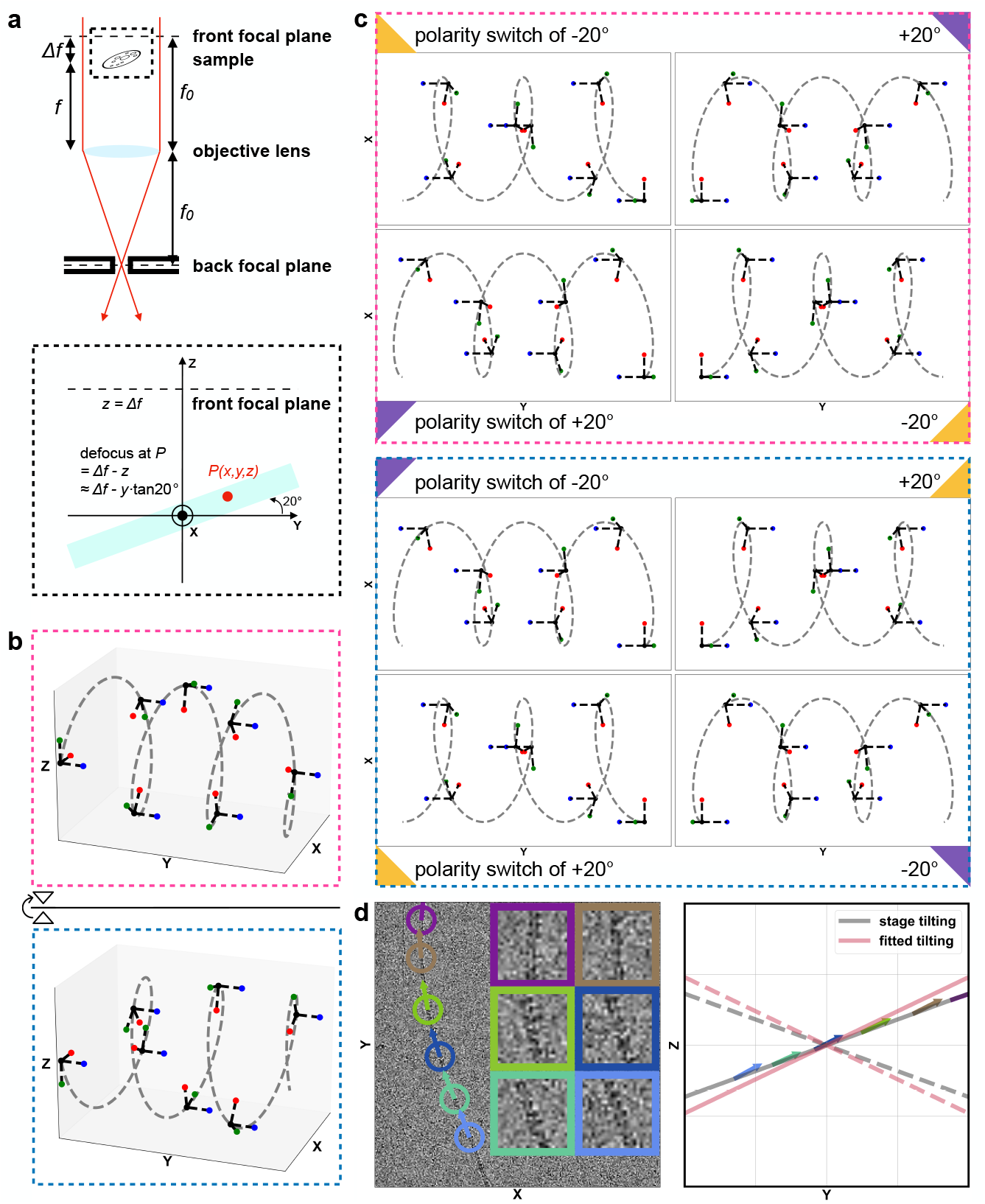
Principle of Ahaha. **a**, The diagram illustrates the optical path of the cryo-EM system with a lens configuration set to underfocused conditions. In this scenario, the sample stage is positioned between the front focal plane of the objective lens and the objective lens itself. *f*_0_ is focal length and Δ*f* is defocus. The region enclosed by the dashed box is enlarged to further illustrate the coordinate system conventions adopted in this study, specifically a right-handed coordinate system where the Z-axis is oriented opposite to the direction of the electron beam. **b**, A pair of enantiomeric helices intended for illustrative purposes, mirrored across the XY-plane. **c**, Z-axis projections of the helices shown in **b**, corresponding to X-axis rotation of 20°, −20°, polarity switch of 20° and polarity switch of −20°. The color of the dashed box, consistent with that in **b**, indicates the identity of the helix being projected. For each projection, the corresponding right-angled equilateral triangles of the same color (purple or yellow) indicate rotational congruence, and their rotational pattern is consistent with the manner in which right-angled equilateral triangles achieve congruence through rotation. **d**, A glycofibril analyzed by Ahaha, with six particles selected from this fibril and their poses depicted. In XY-plane view, the projections of their pose vectors indicate polarity. Meanwhile, in the YZ-plane view, the projections reveal particle tilting, superimposed with stage tilting and planar fitting of defocus values (solid lines), whereas the dashed line represents the mirrored scenario. The congruence between particle tilting and stage tilting indicates that these particles (with such estimated poses) reconstruct the density map with correct absolute hand.

Figure 1b–c help to visualize these two rules. A pair of enantiomeric helices, designed for illustrative purposes, are mirrored across the XY-plane (Figure 1b). Z-axis projections of the enantiomeric helices are shown in Figure 1c, corresponding to X-axis (stage tilting axis) rotation of 20°, 20°, polarity switch of −20° and polarity switch of −20°. The color of the dashed box is consistent between Figure 1b and Figure 1c, serving to distinguish the two helices of the enantiomeric pair. The aforementioned first rule reflects the distinctness between tilting at 20° and −20° within the same colored dashed box (Figure 1c). To facilitate visualization, we attach right-angled equilateral triangles to the projections, where the different colors (yellow and purple) denote the distinctness. The aforementioned second rule corresponds to the identity between the tilt at 20° within one colored dashed box and the tilt at −20° within the other colored dashed box (Figure 1c). This identity is depicted by the matching colors of the attached right-angled equilateral triangles. The polarity switch induces only an in-plane rotation of 180°, and thus aligns with the rotational congruence of the attached right-angled equilateral triangles of the same color.

Therefore, based on the two aforementioned rules, assuming the helices are perpendicular to the tilting axis, in cryo-EM experiments employing a stage tilt of 20°, the detection of projections annotated with purple right-angled equilateral triangles theoretically indicates that the helix encapsulated by the pink dashed box (the upper helix in Figure 1b) is congruent with the applied stage tilting, thereby confirming this helix has the correct absolute hand. Conversely, the identification of projections marked by yellow right-angled equilateral triangles indicates that the helix enclosed within the skyblue dashed box (the lower helix in Figure 1b) is consistent with the stage tilt, thereby confirming that helix has the correct absolute hand.

However, in experiments, the helices do not align perpendicularly with the tilting axis; instead, they intersect the tilting axis randomly. Additionally, they exhibit out-of-plane characteristics, meaning they do not lie perfectly flat or parallel to the thin vitreous ice. Furthermore, glycofibrils are thin, which gives rise to a signal discrepancy between the projections of the tilted enantiomeric pair only at high frequencies. Under such circumstances, relying solely on visual observation of averages resolved through 2D classification is insufficient to determine their absolute hand (Supplementary Figure 1). Thus, 3D refinement coupled with statistical analysis of the pose estimation derived during such refinement, is imperative.

Therefore, the second step in the Ahaha workflow involves 3D refinement of the particles, for obtaining the pose of each particle. The 3D refinement yields a density map that is randomly assigned to one of the the enantiomeric pair. The tilting component of the particle poses estimated during 3D refinement exhibits a sign reversal between the two randomly assigned enantiomers, as will be further elaborated below. Assuming that the signal of the Z-axis projection annotated with purple right-angled equilateral triangles (Figure 1c) is present in the noisy particles, if the random assignment of the absolute hand during 3D refinement results in the upper helix in Figure 1b, the tilt component of the particle poses will be estimated to be 20°. In contrast, if the random assignment of the absolute hand during 3D refinement results in the lower helix in Figure 1b, the particle tilting will be estimated to be −20°. Given that a stage tilting of 20° was employed in cryo-EM data acquisition, this tilt consistency indicates that the upper helix in Figure 1b, output by 3D refinement which estimates particle tilting being 20°, possesses the correct absolute hand. In the case where the signal of the Z-axis projection annotated with yellow (rather than purple) right-angled equilateral triangles (Figure 1c) is present in the noisy particles, vice versa.

Take a glycofibril analyzed by Ahaha as an example, with six particles selected from this fibril and their poses depicted (Figure 1d). In XY-plane view, the projections of their pose vectors indicate polarity. Meanwhile, in the YZ-plane view, the projections indicate particle tilting (obtained in the 3D refinement), superimposed by stage tilting. The congruence between particle tilting and stage tilting indicates that these particles reconstruct the density map with correct absolute hand with such estimated poses. To prevent users of Ahaha from confusing the sign of stage tilting, Ahaha also provides a planar fitting of the defocus values of particles as a double-check (Figure 1d), with defocus estimated by the patch CTF module of cryoSPARC [20].

### 2.2 Correctness validation of Ahaha

The nature-sourced sample, consisting of fresh water collected from a dripping stalagmite in a karst cave (E110.342830°, N25.171496°) was used to validate the correctness of absolute hand determined by Ahaha. This sample was selected for its content of both glycofibrils and bacterial pili with diverse morphologies. Given that bacterial pili are composed of proteins, their absolute hand can also be clearly determined by observing the helical handedness of the *α*-helix [21]. Among the all three bacterial pili measured in this water sample, the absolute hand determined by Ahaha was consistent with that obtained from *α*-helix observations. A detailed analysis is provided as follows.

Density maps with resolutions ranging from 2.9Å to 3.4Å were reconstructed by processing 26,454 mono-tilt cryo-EM micrographs of this water sample (Supplementary Table 1). These micrographs included 12,389 acquired at a 20° tilt and 14,065 acquired at a 30° tilt. As previously stated, Ahaha is to examine whether congruence between particle tilting and stage tilting holds at the level of sign. Consequently, a range of different stage tilts may be employed, provided the sign of the stage tilt remains unchanged. For each of the three bacterial pili, a pair of enantiomeric density maps, which are mirror volumes of each other, was obtained (Figure 2a). Helical symmetry was imposed during 3D reconstruction. The helical parameters of these bacterial pili, which comprise helical rise and twist, had been determined in prior experiments (Table 1).

**Fig. 2.**
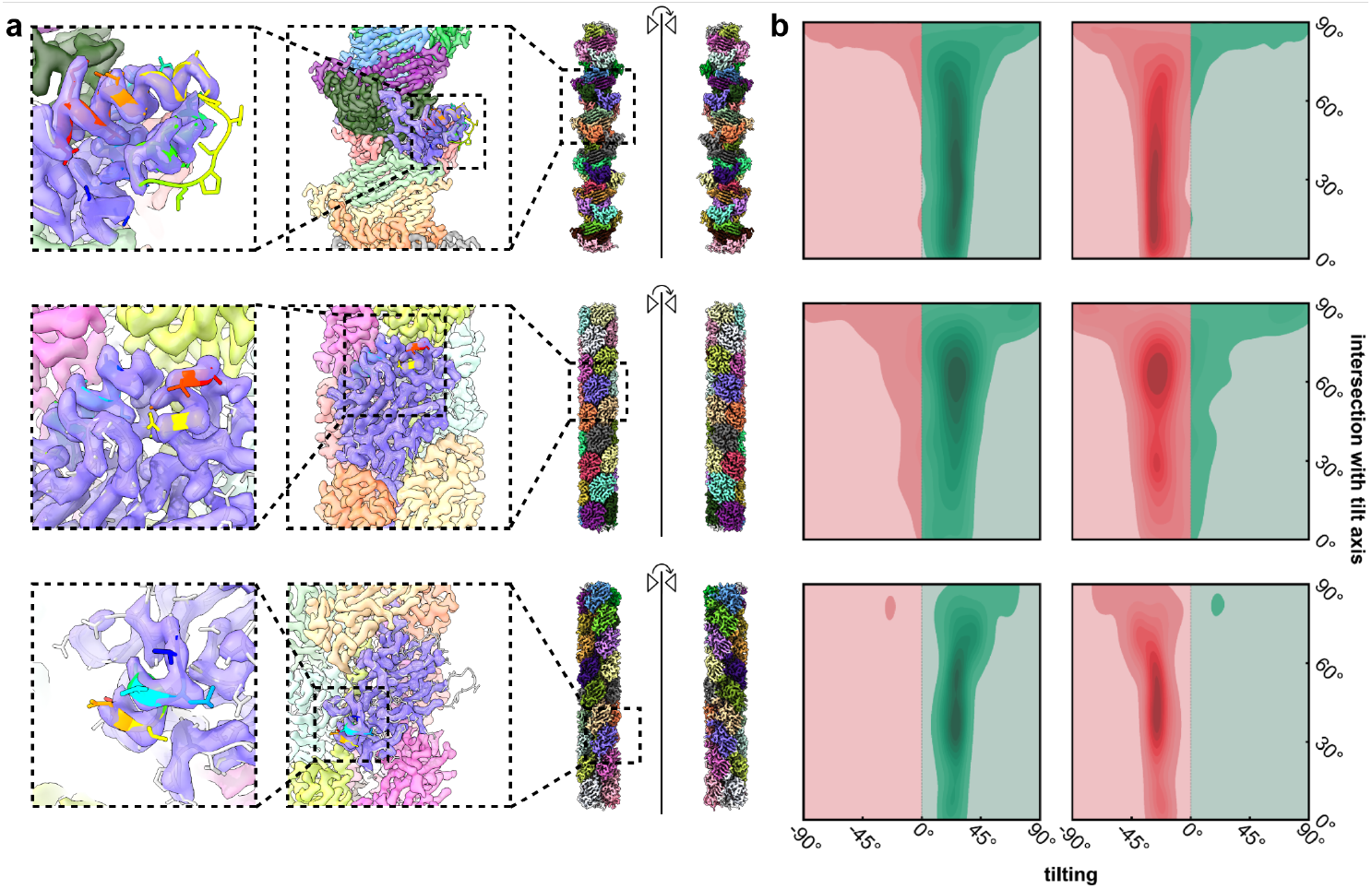
Validation of Ahaha correctness using bacterial pili. The three rows correspond to pilus-like-*α*, pilus-like-*β*, and pilus-like-*γ*, respectively. **a**, The density maps of the enantiomeric pair, shown on the right, are mirror volumes of each other, with different subunit colored distinctly. The left member of the mirror pair exhibits the correct absolute hand, as the *α*-helices are right-handed. The *α*-helical regions within the density maps are shown in an enlarged view, overlaid with atomic models; segments containing *α*-helices are colored according to a spectral gradient (from red to blue), corresponding to the progression from the N-terminus to the C-terminus. **b**, With Ahaha analysis, the tilting of the particles are depicted as heatmaps, plotted against their attributes of intersection with the tilt axis. The left and right heatmaps correspond to the correct-handed and wrong-handed density maps, respectively. Positive tilting values (green) are consistent with stage tilting, whereas negative tilting values (red) are opposite to stage tilting.

Density maps corresponding to the true absolute hand of the enantiomeric pair were identified by observing the helical handedness of the *α*-helix, which is expected to be right-handed. For the density maps with true absolute hand, atomic models were automatically constructed using EModelX [22], with concurrent determination of the amino acid sequence (Supplementary Figure 2-7, Supplementary Table 2). The identities of three bacterial pili (pilus-like-*α*, pilus-like-*β*, and pilus-like-*γ*) were determined via amino acid sequence alignment, with their closest homologs listed in Supplementary Table 2. The built atomic models facilitated the illustration presented in Figure 2a, where atomic models of the *α*-helix-containing segments are colored according to a spectral gradient from red to blue, corresponding to the progression from the N-terminus to the C-terminus.

Ahaha was subsequently utilized to determine the absolute hand of these density maps by analyzing the corresponding poses of the particles that contribute to their reconstruction. For all three bacterial pili, the correctness of Ahaha is validated by the sign-level congruence between the tilting (within a range proximal to 20° and 30°) of particles that reconstruct density maps with the true absolute hand, and the stage tilting (20° and 30°) employed in cryo-EM data acquisition (Figure 2b, green halves of the heatmaps). In contrast, the other members of the enantiomeric pairs, which corresponds to the mirroring volumes of the correct-handed density maps, exhibits respective tilting of their particles within a range proximal to −20° and −30°, differing from the stage tilting by a sign reversal (Figure 2b, red halves of the heatmaps).

### 2.3 Absolute hand measurements of glycofibrils

Given the validated accuracy of absolute hand determination by Ahaha, we selected four glycofibrils derived from the same water sample for absolute hand measurements. The same dataset composed of 26,454 mono-tilt cryo-EM micrographs was processed to obtain density maps of these four glycofibrils, with resolutions ranging from 3.2Å to 3.7Å (Supplementary Table 1). With Ahaha, all four glycofibrils exhibited a clear tilting tendency (Figure 3a). Thus, the correct members of the enantiomeric pairs were thereby determined. Positive particle tilting values (Figure 3a, green halves of the heatmaps) are consistent with stage tilting, whereas negative particle tilting values (Figure 3a, red halves of the heatmaps) run counter to stage tilting; accordingly, the left density map displays the correct absolute hand. The rise and twist of the density maps with the correct absolute hand are illustrated in Figure 3b.

**Fig. 3.**
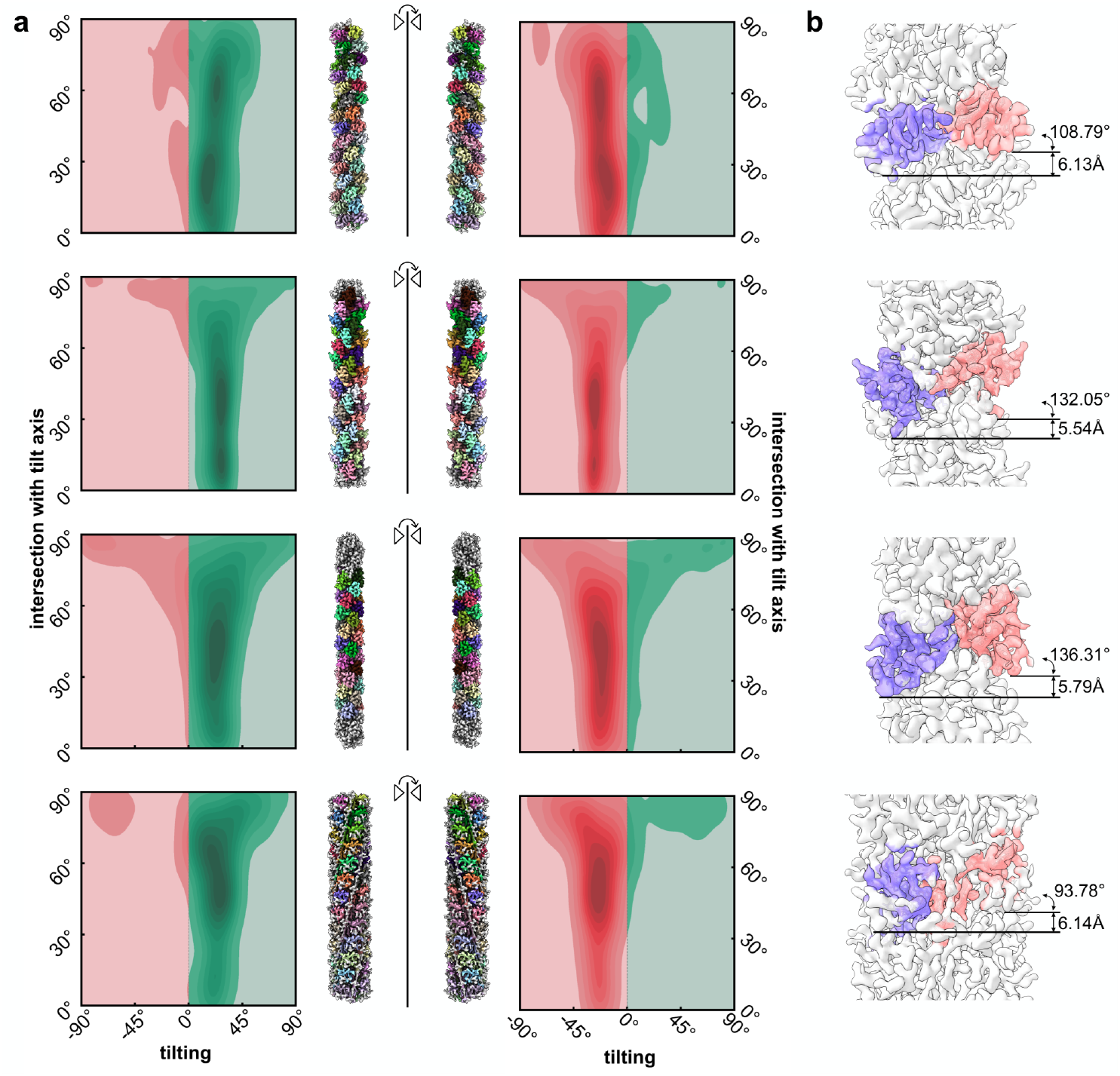
Absolute hand measurements of glycofibrils. The four rows corresponds to TLP-2a, TLP-2f, TLP-2g and TLP-2h, respectively. **a**, The density maps of the enantiomeric pairs are depicted as volumes that mirror each another. Regions of the density that can be subsequently modeled are colored, with coloring distinguishing the asymmetric unit. Through Ahaha analysis, the tilting of particles is visualized as heatmaps, which are plotted against their attributes of intersection with the tilt axis. These heatmaps are positioned alongside the corresponding density map within the enantiomeric pair. Positive tilting values (green) are consistent with stage tilting, whereas negative tilting values (red) are counter to stage tilting; consequently, the left density map exhibits the correct absolute hand. **b**, For the correct-handed glycofibril density map, helical rise and twist between the two adjacent asymmetric units, colored purple and red, are labeled herein.

### 2.4 Atomic models of glycofibrils

Once the absolute hand was determined, the high-resolution density maps (2.8Å to 3.1Å) of four glycofibrils, acquired by processing non-tilted micrographs (Supplementary Table 1) from preceding experiments, underwent model building. The decision to flip the maps or not was determined by their consistency with the corresponding low-resolution density maps (3.2Å to 3.7Å, correct absolute hand) obtained from mono-tilt micrographs and Ahaha analysis (Supplementary Table 1, Figure 3). Upon model building, we observed that all of them are glycan-coated linear chains composed of dipeptide repeats (Figure 4), despite substantial differences in their overall morphology (Supplementary Figure 8). Their distinct morphological characteristics arise not only from variations in the mode of O-glycosylation but also from differences in the coated polysaccharides. A detailed analysis is provided as follows.

**Fig. 4.**
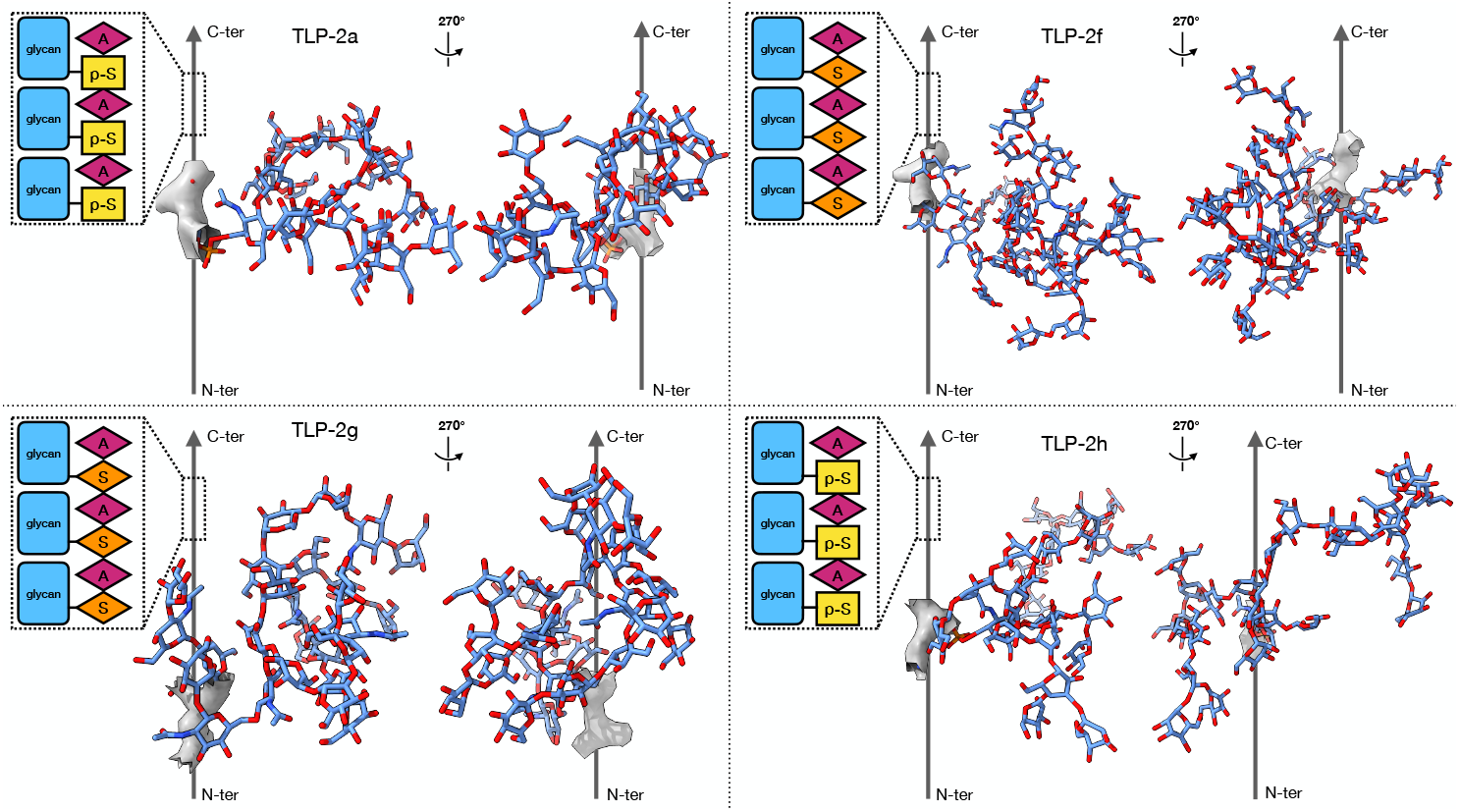
Atomic models of the glycofibrils. The single glycan linked to Ser or p-Ser is depicted for each TLP-2 subtype. The gray arrow indicates the linear chains composed of dipeptide repeats, though N-ter/C-ter of them was inferred by observing the oxygen atoms in the carboxyl groups of the amino acids. Higher-resolution density maps are required for definitive confirmation. The diagram enclosed within the dashed box illustrates both the composition of the linear chains and the O-glycosylation site. Owing to the limited resolution, the types of sugar molecules incorporated into the model constitute no more than our most reasonable approximations. In instances where the density lacks a clear propensity for a particular sugar type, mannose was chosen as the default.

For TLP-2f and TLP-2g, the glycosylation mode can be immediately identified as O-glycosylation at Ser/Thr residues [23], as this pattern provides the best fit to the corresponding density map (Supplementary Figure 9). For TLP-2a and TLP-2h, the N-glycosylated [24] Asn residues are not supported by the density map, and the distance between the sugar moiety and the amino acid is also inconsistent with O-glycosylation at Ser/Thr (Supplementary Figure 9). Therefore, we speculate that TLP-2a and TLP-2h represent another type of glycosylation modification. Phosphoglycosylation involves the attachment of a phosphoglycan to Ser/Thr residues [25, 26]. Notably, when we modeled a phosphate group into the corresponding density, it fit perfectly (Supplementary Figure 9). Therefore, TLP-2a and TLP-2h may represent phosphoglycosylated fibrils. The four glycofibrils also exhibit significant differences in their coated polysaccharides. In Figure 4, the single glycan linked to Ser or p-Ser is depicted for each TLP-2 subtype. The glycans pack against one another via O-glycosylation linking to every Ser or p-Ser, forming a helical assembly.

However, due to the limited resolution (2.8Å to 3.1Å), the N-ter/C-ter of the core linear chains composed of dipeptide repeats was inferred by observing the oxygen atoms in the carboxyl groups of the amino acids. Higher-resolution density maps are required for definitive confirmation. Moreover, also owing to the limited resolution, the types of sugar molecules modeled represent merely our best approximations. In cases where the density does not exhibit a tendency toward a specific sugar type, mannose was selected as the default.

## 3 Discussion

Ahaha is a straightforward and effective method for determining the absolute hand of helical assemblies. It relies solely on the analysis of mono-tilt micrographs. Given the current cryo-EM data acquisition strategy, mono-tilt micrographs can be acquired in a high-throughput manner. The limitation of Ahaha is that it has a lower bound for fibril abundance. Specifically, extremely rare glycofibrils, which are at the detection limit of CryoSeek, may not provide sufficient signal to enable absolute hand determination via Ahaha. Another potential limitation of Ahaha is that it may be ineffective when the diameter of the glycofibril is extremely small. However, the minimum diameter of glycofibrils for which Ahaha can reliably determine the absolute hand remains to be investigated.

In this study, we built atomic models of four glycofibrils, based on their absolute hand determined by Ahaha. They are sourced from just one single water sample, specifically the water droplets from a stalagmite within a karst cave. As discussed in our previous studies [6, 8], upon examining micrographs of nature-sourced samples (e.g., natural water samples), substantial components, including a large number of fibrils with diverse morphologies, can be observed with naked eyes. Among the glycofibrils that constitute these fibrils, their taxonomy is of particular interest. In our previous study, we reported TLP-4 [7], a glycan-coated linear chain composed of tetrapeptide repeats with a rise of 12.4Å. The four glycofibrils reported in this study, whose protein moieties consist of dipeptide repeats, exhibit rises of 6.13Å, 5.54Å, 5.79Å, and 6.14Å (Table 1), values that are close to half of the rise observed for TLP-4. This may suggest the prevalence of glycan-coated linear chains with repeated peptides, albeit with diverse glycosylation patterns. In other words, glycofibrils characterized by a linear chain as the central axis (as the protein moiety), coated with various types of O-glycosylation, and associated with distinct polysaccharides, may form a family within the potential taxonomy of glycofibrils. Furthermore, the number of amino acids per repeat may constrain the helical rise in the assembled structure, whereas the twist angle can vary. Although we have thus far only observed dipeptide and tetrapeptide repeats as the linear chain, hexapeptide, octapeptide, or even odd-numbered-peptide may also exist as repeating units.

Despite their biological significance, glycans remain less well understood compared to proteins and nucleic acids, primarily due to their high diversity and complex architectures, including encompassing varied compositions, configurations, and linkage patterns. Most characterized glycan structures to date are relatively simple, typically consisting of a small number of glycosyl units attached to the extracellular loops of membrane proteins or secreted proteins. Against this backdrop, CryoSeek has opened a novel avenue for glycan research by enabling the high-throughput structural determination of glycofibrils from natural environments, expanding the scope of investigable glycan structures. However, prior to the present study, absolute hand determination for glycofibrils remained unachievable. Ahaha fulfills this final critical step in cryo-EM image processing workflows, thereby advancing glycan research through their structural investigation.

## Methods

### Pretreatment of fresh water collected from the cave

The water sample utilized in this study was collected on 11 May 2025 from a dripping stalagmite within a karst cave located in Yanshan district, Guilin, China (E110.342830°, N25.171496°). Upon sample preparation, 5 liters of collected water were subjected to filtration with pore size 0.22*µ*m (JinTeng). The filtrate underwent concentration to a final volume of 200 mL via a tangential flow filtration system (Merck), utilizing a 0.11 m^2^ membrane (Biomax) with a 50-kDa cutoff, under a constant transmembrane pressure of 0.7 bar at ambient temperature. The solution was further concentrated to approximately 50*µ*L using Centricons (Millipore) with a 100-kD cutoff.

### Cryo-EM sample preparation and data acquisition

Cryosamples were prepared using a Vitrobot Mark IV (Thermo Fisher Scientific). Aliquots of 4*µ*L of concentrated aqueous solution were deposited onto glow-discharged holey carbon grids (Au 300 mesh, R1.2*/*1.3, Quantifoil). Following a blotting duration of 6 seconds, the grids were plunge-frozen in liquid ethane that had been precooled with liquid nitrogen.

Cryo-EM data acquisition were conducted with two Krios G4 300 kV transmission electron microscopes, each outfitted with a Falcon4i direct electron detector and a Selectris X energy filter (with a slit width of 10 eV). Movies were acquired in EER mode with a total does of 50*e*^−^/*Å*^2^ with EPU. The magnification was set to *×*130, 000, corresponding to pixel sizes of 0.927Åand 0.936Åfor each microscope, respectively, with a defocus range from 1.2 to 1.6*µ*m. Of a total of 26,454 mono-tilt movies, 12,389 and 14,065 were recorded with the stage tilted at 20° and 30°, respectively. Additionally, 22,272 movies were acquired under non-tilted conditions.

### Image processing and 3D reconstruction of density maps

Following motion correction and CTF estimation performed using cryoSPARC [20], approximately 12 million and 15 million particles were selected via the filament tracer module from 22, 272 non-tilted movies and 26, 454 mono-tilt movies, respectively.

Following multiple rounds of 2D classification, particles suitable for high-resolution reconstruction were selected (metadata regarding the particles and their reconstruction can be found in Supplementary Table 1). Helical symmetry was imposed during 3D reconstruction, with helical parameters determined either via helical indexing in Fourier space using AI-HEAD or in real space using HI3D [27]. The aforementioned 2D classification and 3D reconstruction were also performed using cryoSPARC [20].

Enantiomeric pairs of density maps can be generated through the following steps: first, the initial refinement volume is flipped; subsequently, two mirror-symmetric twist angles are imposed during symmetry enforcement in the refinement process, with *θ* applied to the initial volume and 360° − *θ* applied to the flipped initial volume, where *θ* represents the twist in the helical parameters.

### Absolute hand measurement by Ahaha

Following image processing of mono-tilt micrographs, high-resolution density maps were obtained, alongside the corresponding particles and their poses. The poses of the particles, together with their associated filament tracing data, serve as input for the Ahaha analysis. Ahaha initially filters out filaments composed of fewer than 10 particles. Furthermore, to eliminate noise introduced by inaccuracies in classification and orientation determination [13, 28, 29], which may induce contradictory particle alignments within individual filaments, Ahaha employs an entropy-based filtering criterion defined as follows:

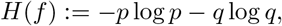

where *p* represents the fraction of particles with helical refinement-derived orientations consistent with the filament’s axial direction, and *q* = 1 − *p* denotes the inconsistent fraction. This entropy metric quantifies the reliability of polarity estimation for each filament, with higher values indicating greater disorder in the refined orientations. Ahaha applies a stringent entropy threshold (set to 0.01 in our experiments) to exclude filaments with poorly determined polarity. Remarkably, even with this conservative threshold, all fibril types studied retained at least 45 high-quality filaments, demonstrating the consistent presence of well-ordered structural elements in our samples suitable for robust absolute hand determination.

Ahaha calculates the intersection between the pose of each particle and the tilt axis. This is because particles aligned parallel to the tilt axis yield minimal tilt-related information, rendering them less informative for absolute hand determination. By integrating particle tilting and the intersection with the tilt axis, heatmap visualizations were constructed. Particles that are consistent with the applied stage tilt are represented in green, indicating a positive absolute hand, whereas those exhibiting an opposing tilt are displayed in red, signifying a negative absolute hand.

### Model building

For pilus-like-*α*, pilus-like-*β* and pilus-like-*γ*, the reconstructed density map was directly fed into EModelX(w/o seq), which automatically generated initial models of the helical assemblies. Taking pilus-like-*α* as an example, a zoomed-in view of an individual protomer reveals that the auto-built model (Supplementary Figure 2b, skyblue trace) aligns well with the map. Concomitantly, EModelX predicted the underlying protein sequence for the protomer, which yielded a high-confidence match to UniProt entry A0A257CJ09 (BLAST [30] E-value = 2.2 *×* 10^−56^). The AlphaFold-predicted structure [31, 32] of A0A257CJ09 (Supplementary Figure 2a, gray trace) showed strong structural similarity to the auto-built model (TM-score [33] = 0.844). Both sequence and structural alignments support the assignment of A0A257CJ09 as the corresponding sequence of pilus-like-*α*. Therefore, we reran EModelX with the guidance of A0A257CJ09 to automatically build a homomeric assembly. Finally, manual refinement was performed with Coot [34], during which the amino acid identity might be adjusted according to the map density (Supplementary Figure 2b, darkorange trace). The refined model exhibited favorable alignment with the density map (Supplementary Figure 2c). Analogous procedures were performed for pilus-like-*β* and pilus-like-*γ* (Supplementary Figure 3-4).

For TLP-2a, TLP-2f, TLP-2g and TLP-2h, atomic models were built manually with Coot [34].

## Code availability

The online service of Ahaha can be accessed via https://cryoseek.org/ahaha.

## Acknowledgements

We thank Dr. Gang Fu and Xiaosong Pei for their technical support during cryo-EM image acquisition. We also thank the Structural Biology Core Facility at the Bio-Tech Center of Shenzhen Medical Academy of Research and Translation (SMART) for their technical assistance. Additionally, we acknowledge the support in data analysis provided by the Computing Labware for Electronmicroscopy Visualization and Experimental Research (CLEVER) at SMART. We are grateful to Dr. Li Xu for her valuable guidance on atomic model building. We thank Dr. Juanhao Huang for the fruitful discussions we had with him. We thank Fei Zhang for her support in collecting water samples. We would like to express our sincere gratitude to Prof. Yifan Cheng for his insightful suggestions contributed during the development of this manuscript. This work was supported by SMART (to M.H.), Beijing Frontier Research Center for Biological Structure (Tsinghua University) (to M.H.), Beijing Advanced Innovation Center for Structural Biology (Tsinghua University) (to M.H.), National Natural Science Foundation of China (32371254 and 32171190 to J.W., 92478205 to N.Y.).

## Contribution

M.H., J.W., and N.Y. initiated the project. M.H. formulated the principle of Ahaha, while Q.Z. carried out its implementation. L.Q. collected the water sample, while L.Q. and Y.Z. prepared the sample and conducted cryo-EM data acquisition. M.H. performed cryo-EM image processing, while Q.Z. conducted the Ahaha analysis. J.W. and S.C. built atomic models. T.W., Z.L., and J.W. analyzed the atomic models. Z.L. and T.W. identified O-glycosylation at the p-Ser sites in TLP-2a and TLP-2h. M.H., Q.Z., J.W., S.C., Z.L., and N.Y. wrote the manuscript.

