## Supplementary material for "Absolute hand determination of glycofibrils from natural sources in cryo-EM": SI

### Supplementary Note I

We employ the term “absolute hand” rather than “handedness” due to the following consideration: within the context of helical assembly, the conventional designation defines a twist ranging from $0^\circ$ to $180^\circ$ as right-handed, whereas a twist ranging from $180^\circ$ to $360^\circ$ (equivalent to $-180^\circ$ to $0^\circ$) is referred to as left-handed. This is conventionally termed the handedness of the helical assembly.

Due to the intrinsic loss of absolute hand information in cryo-EM transmission imaging, 3D reconstruction generates a density map that is randomly assigned to one member of an enantiomeric pair. In the case of helical assembly, the two mirroring helices correspond to left-handed and right-handed configurations, respectively. The absolute hand determination method described in this manuscript discriminates between the two enantiomers to identify the correct one, rather than directly establishing whether the true twist lies within the range of $0^\circ$ to $180^\circ$ or $180^\circ$ to $360^\circ$. However, once the correct enantiomer is identified, the handedness of the helix (i.e., left-handed or right-handed) is thereby determined.

### Supplementary Note II

Nature-sourced glycofibrils exhibit high heterogeneity in samples. When using tilt-pair imaging to determine their absolute hand, the identity of individual particles remains unknown until high-resolution reconstruction is achieved. Consequently, high-resolution reconstruction must be performed alongside with tilt-pair imaging. This requirement necessitates a massive number of tilt-pair micrographs, if not for all micrographs in the dataset (typically over ten thousands micrographs). Given the time-consuming nature of stage tilting, acquiring such a large tilt-pair dataset is impractical for experiments. Finally, nowadays, the majority of researchers in the single-particle cryo-EM field are unfamiliar with image processing tools for tilt-pair analysis [1–4], particularly because tilt-pair alignment requires manual intervention and technical expertise. If a massive number of tilt-pairs need to be aligned, this process becomes highly labor-intensive.

**Supplementary Table 1**: Reconstruction metadata of bacteria pili and glycofibrils in the studied water sample

| fibril | number of particles | resolution | number of particles | resolution (enantiomeric pair) |
|---|---|---|---|---|
| | non-tilted | | tilted | |
| pilus-like-$\alpha$ | 46,644 | 3.07Å | 138,530 | 2.94Å/2.92Å |
| pilus-like-$\beta$ | 12,518 | 3.37Å | 34,504 | 3.18Å/3.14Å |
| pilus-like-$\gamma$ | 2,279 | 3.66Å | 6,859 | 3.42Å/3.45Å |
| TLP-2a | 8,688 | 3.10Å | 11,605 | 3.67Å/3.69Å |
| TLP-2f | 16,645 | 2.91Å | 27,262 | 3.28Å/3.29Å |
| TLP-2g | 38,390 | 2.79Å | 66,630 | 3.15Å/3.18Å |
| TLP-2h | 5,082 | 2.80Å | 5,613 | 3.64Å/3.70Å |

**Supplementary Table 2**: Identification of the amino acid sequences of bacterial pili

| pilus | description of the closest homolog | UniProt ID of the closest homolog | AlphaFold database ID of the closest homolog | Sequence alignment with the closest homolog |
|---|---|---|---|---|
| pilus-like-$\alpha$ | DUF6160 domain-containing protein | A0A257CJ09 | AF-A0A257CJ09-F1-v4 | Supplementary Figure 5 |
| pilus-like-$\beta$ | Type IV pilin structural subunit | N8QG80 | AF-N8QG80-F1-v4 | Supplementary Figure 6 |
| pilus-like-$\gamma$ | Type IV pilus assembly protein PilA | A0A0D7KGF4 | AF-A0A0D7KGF4-F1-v4 | Supplementary Figure 7 |

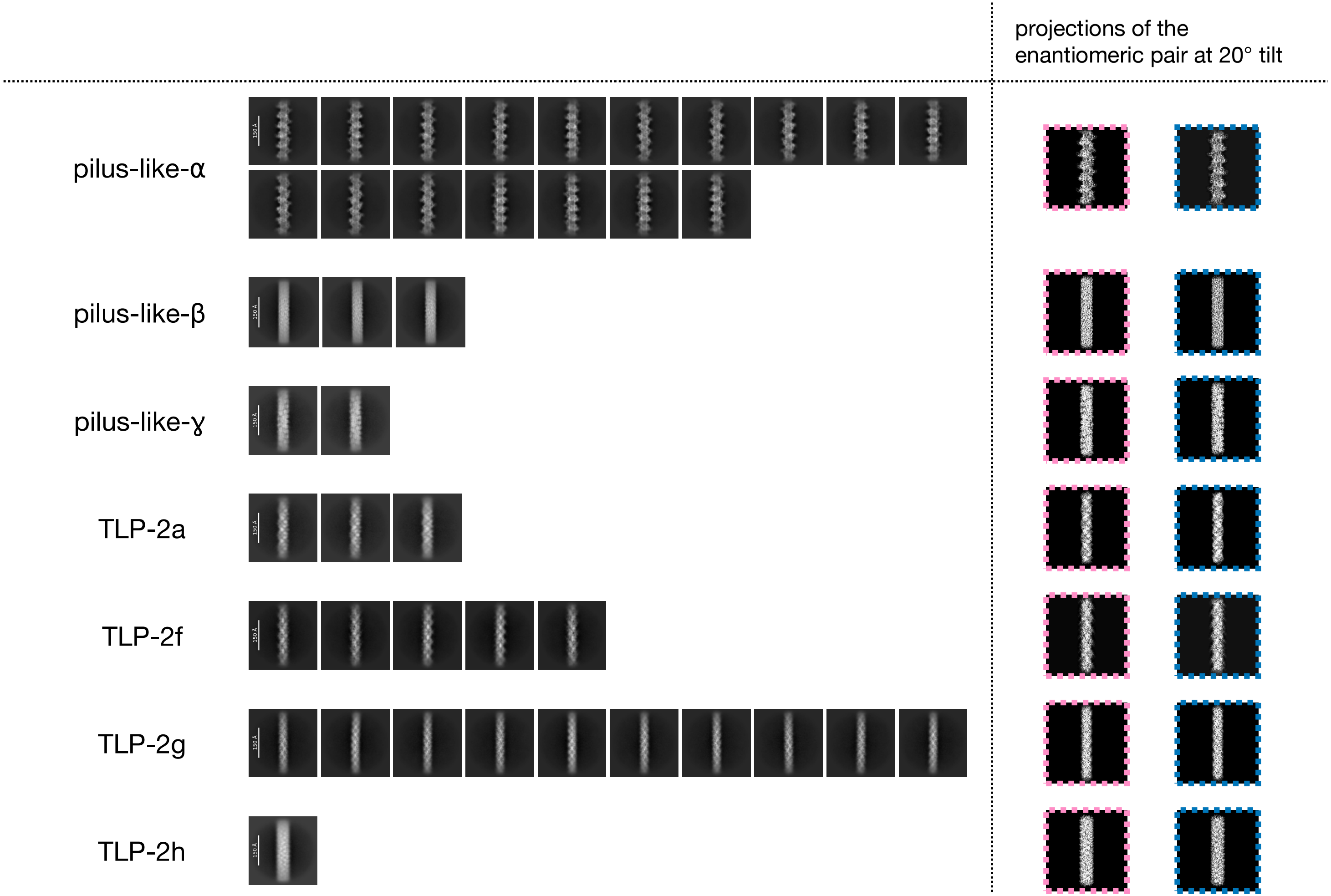


**Supplementary Figure 1**: **The 2D classification averages of fibrils obtained from single-tilt micrographs in this study, alongside projections of the enantiomeric reference density map pairs at a tilting angle of 20°.** The two projections derived from the enantiomeric pair are distinguished by colored dashed boxes.

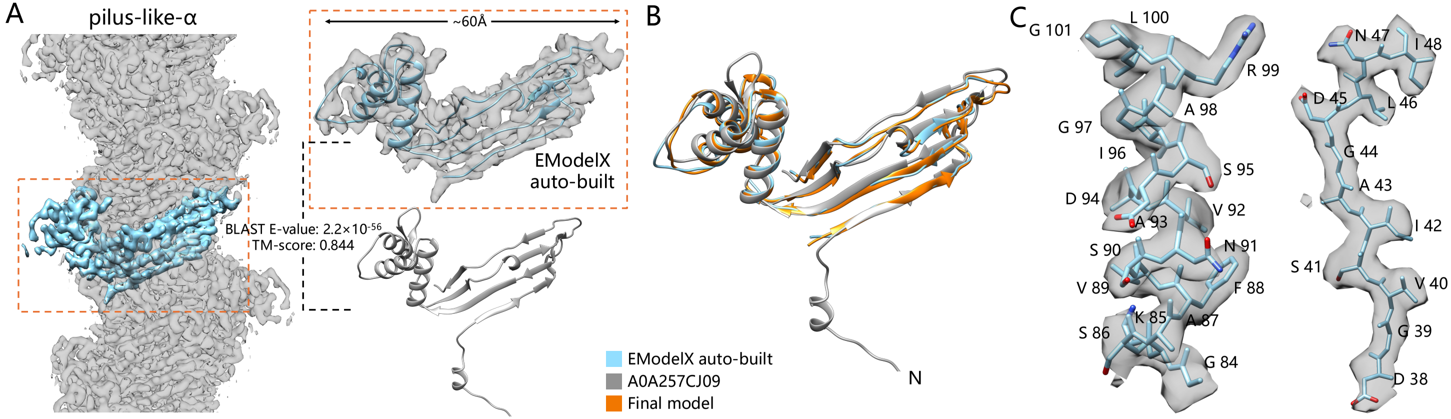


**Supplementary Figure 2**: **Automatic model building of pilus-like-$\alpha$ by EModelX. a**, EModelX initially conducted automatic model building on the selected helical unit (cyan) derived from the cryo-EM density map of pilus-like-$\alpha$, employing an unknown sequence mode. The closest homolog in UniProt (gray) was identified via BLAST using the amino acid sequence predicted in the preceding step. **b**, The final model (darkorange) was obtained through manual fine-tuning based on the cryo-EM density map, with the automatically built model and the closest homolog model serving as the starting points. The automatically built model, the closest homolog, and the final model were aligned for visualization purposes. **c**, Selected segments of the final pilus-like-$\alpha$ model were aligned with their corresponding cryo-EM density maps to assess fitting accuracy.

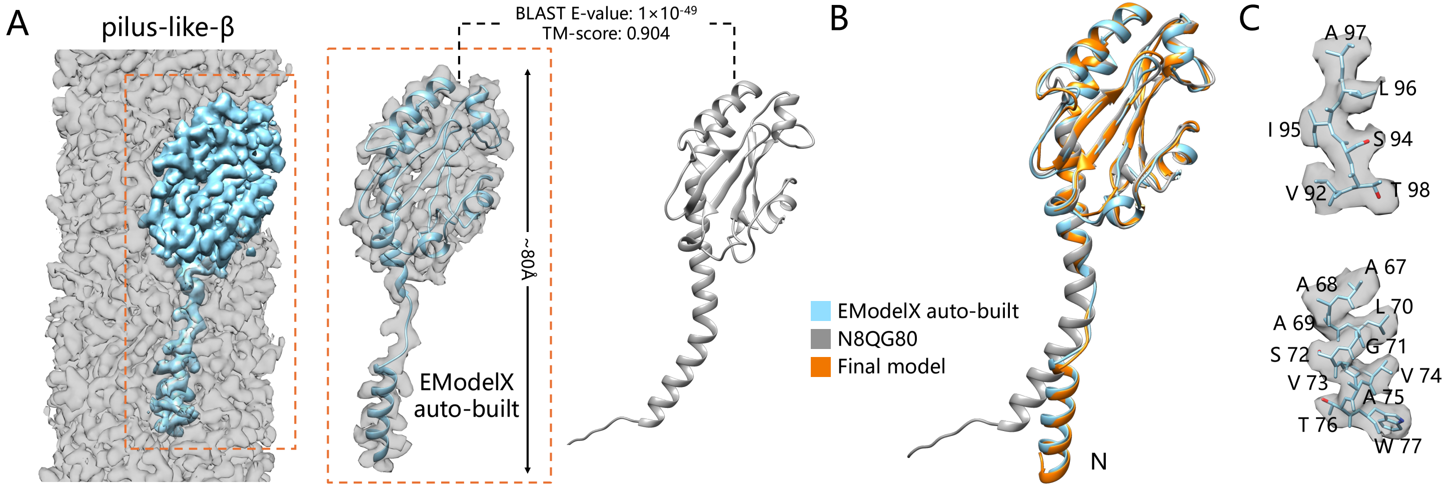


**Supplementary Figure 3**: **Automatic model building of pilus-like-$\beta$ by EModelX.** This figure follows the same structure and content as the caption for pilus-like-$\alpha$ (Supplementary Figure 2), with the only difference being that the protein here is pilus-like-$\beta$ instead of pilus-like-$\alpha$.

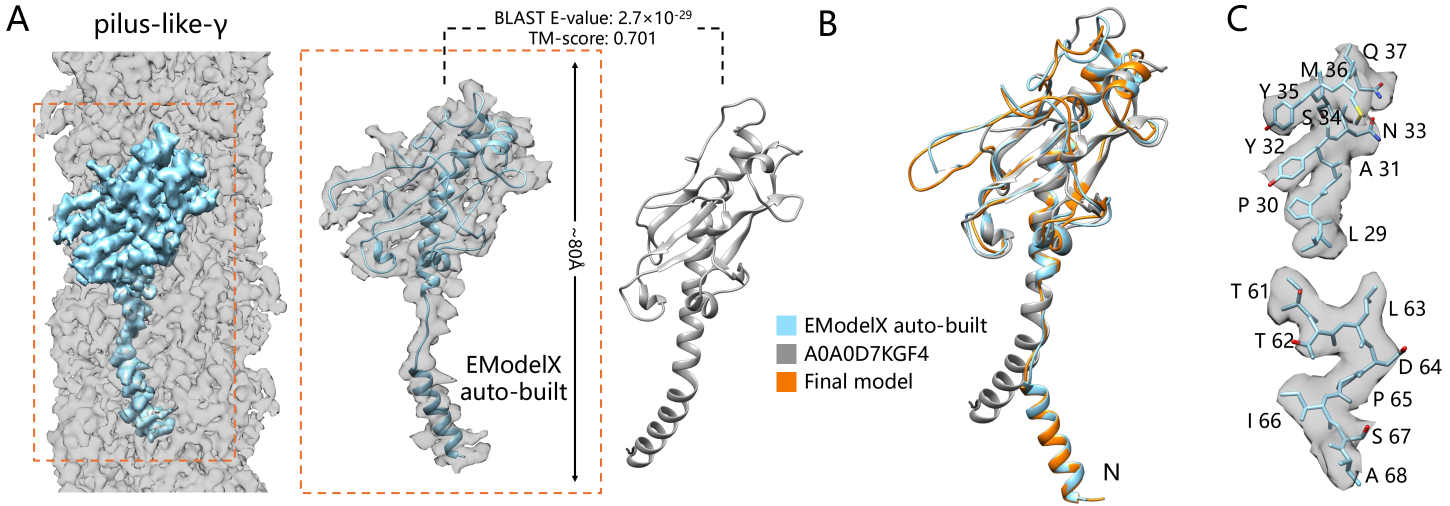


**Supplementary Figure 4**: **Automatic model building of pilus-like-$\gamma$ by EModelX.** This figure follows the same structure and content as the caption for pilus-like-$\alpha$ (Supplementary Figure 2), with the only difference being that the protein here is pilus-like-$\gamma$ instead of pilus-like-$\alpha$.

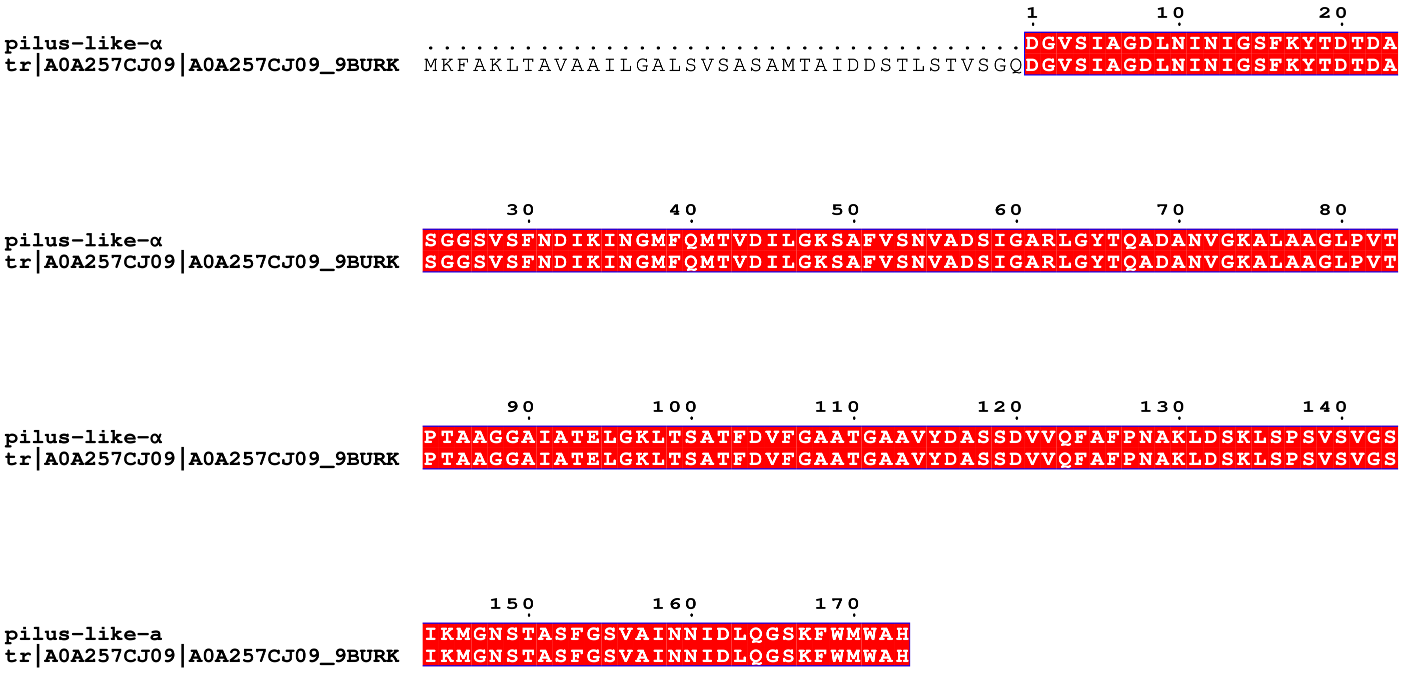


**Supplementary Figure 5**: Sequence alignment of pilus-like-$\alpha$ with its closest homolog (UniProt A0A257CJ09).

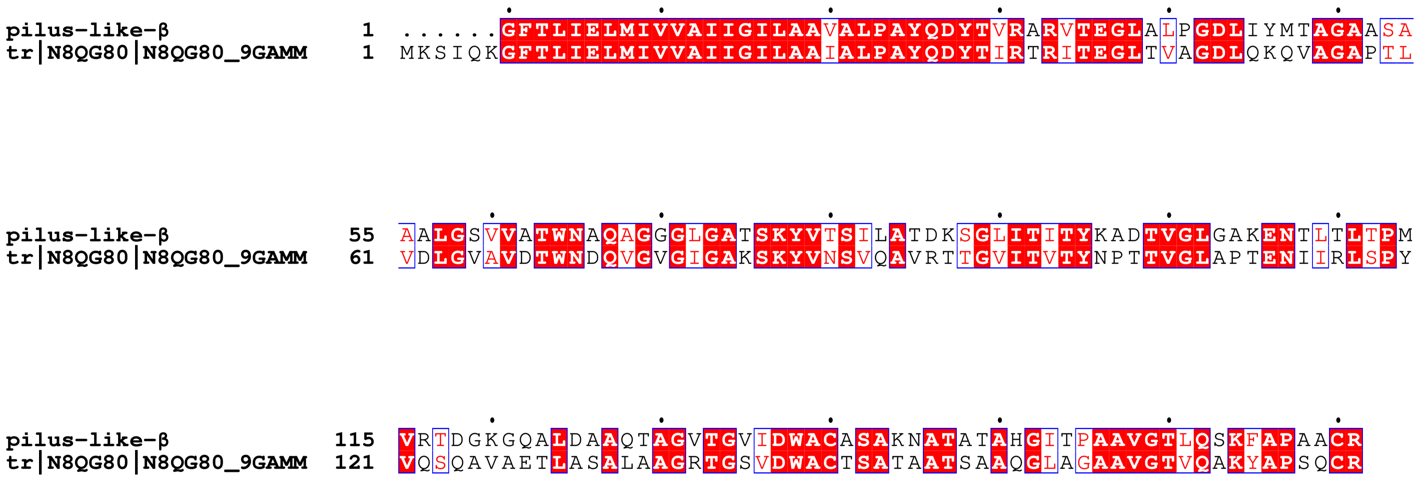


**Supplementary Figure 6**: Sequence alignment of pilus-like-$\beta$ with its closest homolog (UniProt N8QG80).

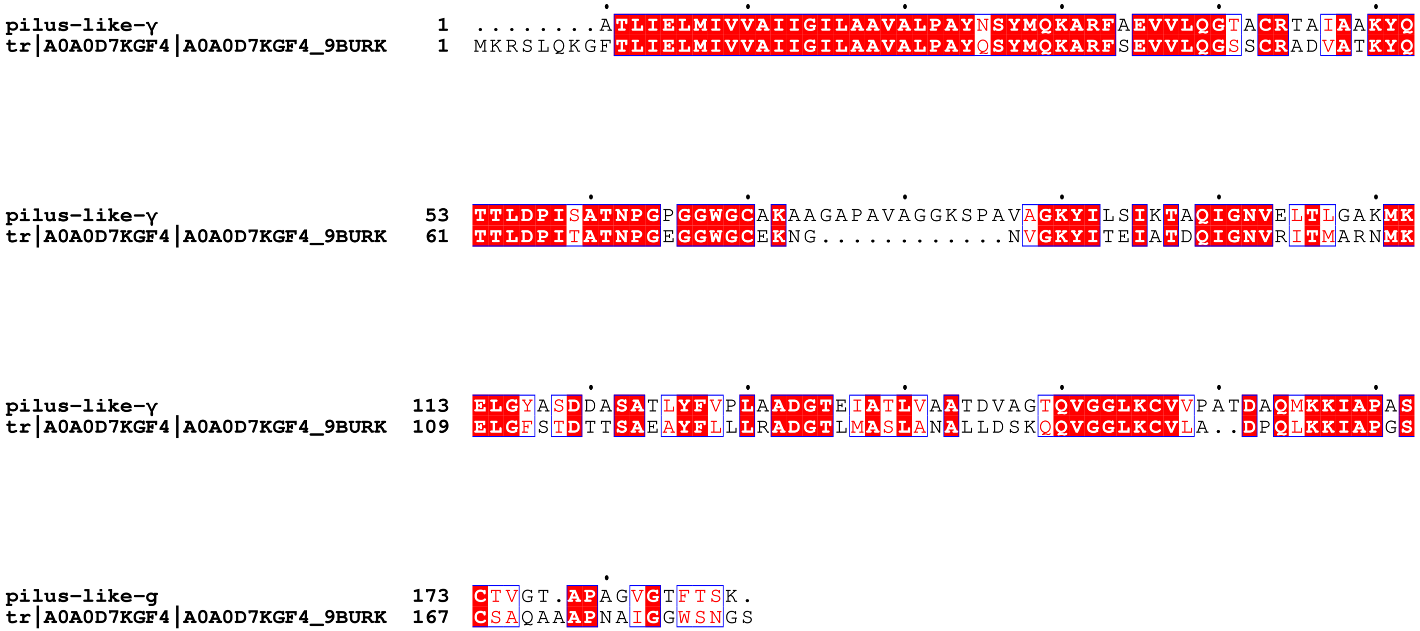


**Supplementary Figure 7**: Sequence alignment of pilus-like-$\gamma$ with its closest homolog (UniProt A0A0D7KGF4).

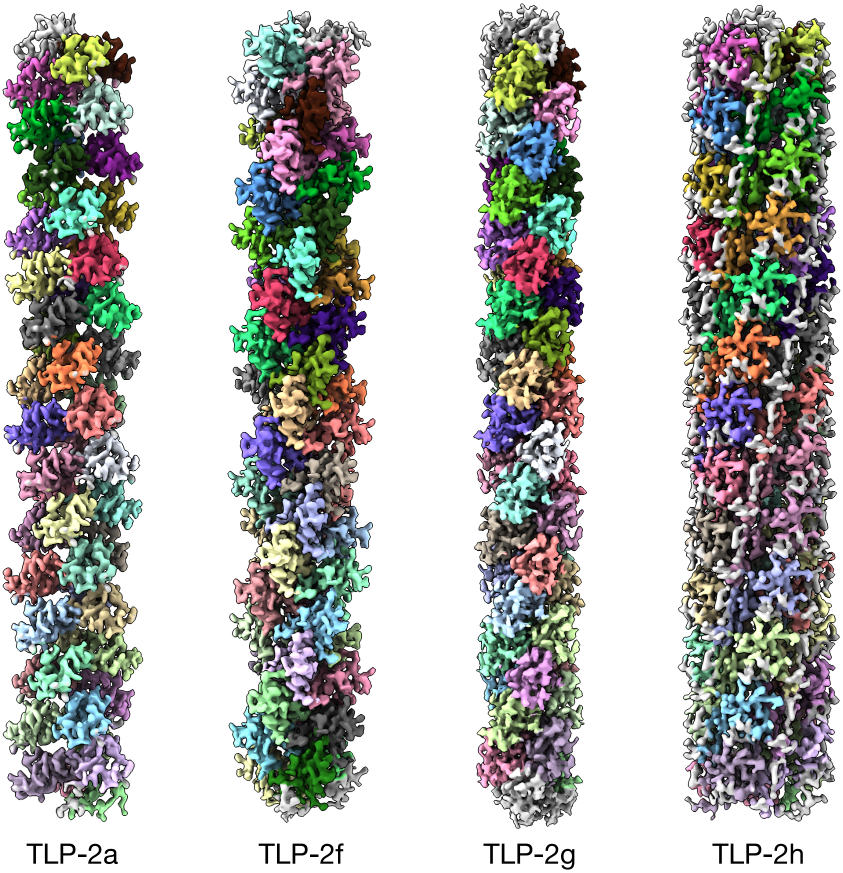


**Supplementary Figure 8**: **Morphologies of TLP-2 subtypes.** Coloration serves to distinguish the asymmetric unit, while regions of density that are ambiguous for model building remain gray.

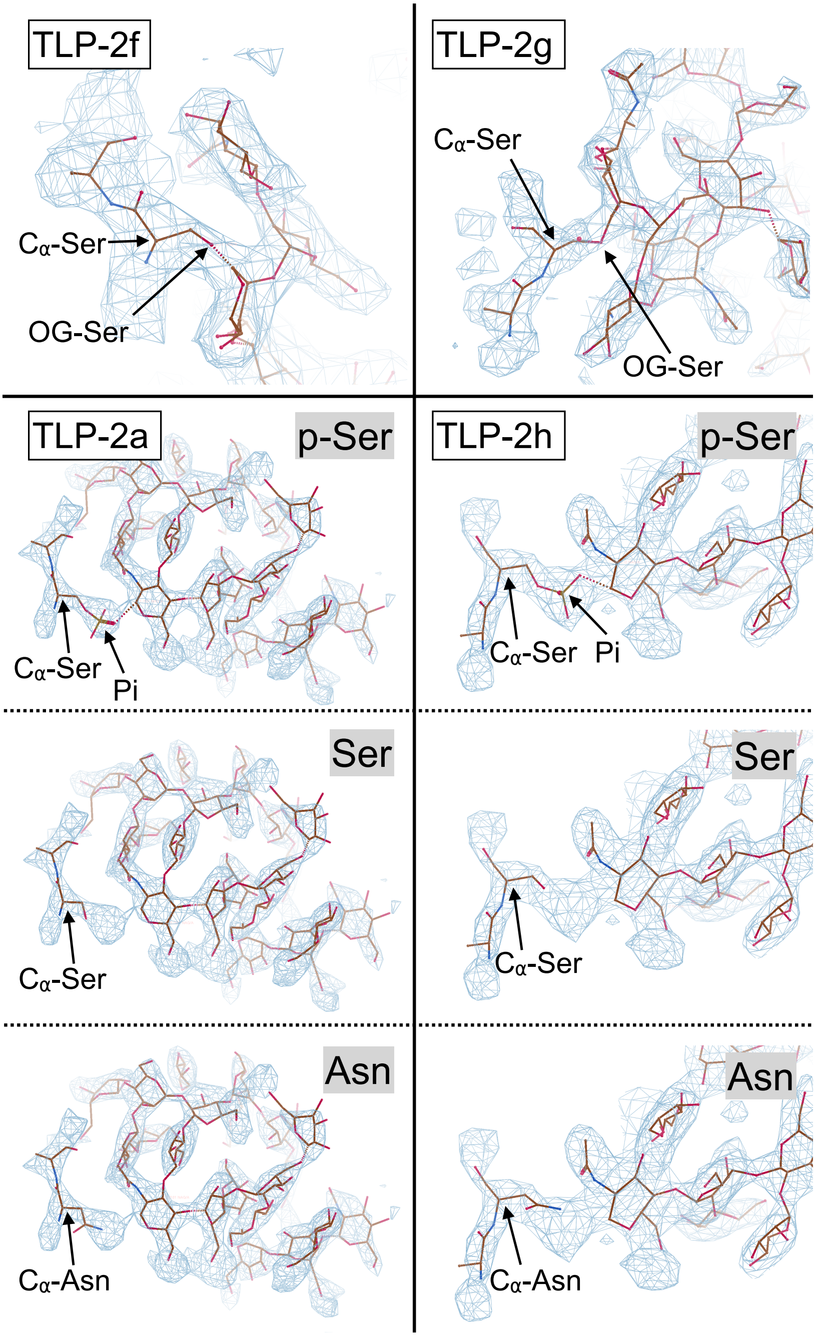


**Supplementary Figure 9**: **Characterization of glycosylation type from density.** In TLP-2f and TLP-2g, the glycan-Ser aligns with the density map. In contrast, in TLP-2a and TLP-2h, the glycan-p-Ser is consistent with the density map, whereas neither glycan-Ser nor glycan-Asn shows such alignment.
